# Characterization of second-order mixing effects in reconstructed cross-spectra of random neural fields

**DOI:** 10.1101/2022.01.19.476889

**Authors:** Rikkert Hindriks

## Abstract

Functional connectivity in electroencephalography (EEG) and magnetoencephalography (MEG) data is commonly assessed by using measures that are insensitive to instantaneously interacting sources and as such would not give rise to false positive interactions caused by instantaneous mixing of true source signals (first-order mixing). Recent studies, however, have drawn attention to the fact that such measures are still susceptible to instantaneous mixing from *lagged* sources (i.e. second-order mixing) and that this can lead to a large number of false positive interactions. In this study we relate first- and second-order mixing effects on the cross-spectra of reconstructed source activity to the properties of the resolution operators that are used for the reconstruction. We derive two identities that relate first- and second-order mixing effects to the transformation properties of measurement and source configurations and exploit them to establish several basic properties of signal mixing. First, we provide a characterization of the configurations that are maximally and minimally sensitive to second-order mixing. It turns out that second-order mixing effects are maximal when the measurement locations are far apart and the sources coincide with the measurement locations. Second, we provide a description of second-order mixing effects in the vicinity of the measurement locations in terms of the local geometry of the point-spread functions of the resolution operator. Third, we derive a version of Lagrange’s identity for cross-talk functions that establishes the existence of a trade-off between the magnitude of first- and second-order mixing effects. It also shows that, whereas the magnitude of first-order mixing is determined by the inner product of cross-talk functions, the magnitude of second-order mixing is determined by a generalized cross-product of cross-talk functions (the wedge product) which leads to an intuitive geometric understanding of the trade-off. All results are derived within the general framework of random neural fields on cortical manifolds.

## 1 Introduction

There are three basic ways in which spurious interactions between two measurement locations can arise in reconstructed source activity, namely through instantaneous linear mixing of incoherent, instantaneously coherent, or lagged coherent cortical activity. In the literature, different names have been used to describe these effects. In [2], the first two ways have been jointly referred to as *first-order artifacts* and the second way as *second-order artifacts*, in [19], the first way is referred to as *artificial synchrony* and the second and third ways are jointly referred to as *spurious synchrony*, and in [18], the third way is referred to as *ghost interactions*. For the analysis in the current study, we find it useful to distinguish all three ways and will refer to them as zeroth-, first-, and second-order mixing effects, respectively. Thus, in this terminology, zeroth-, first-, and second-order effects refer to the effects of instantaneous linear mixing of incoherent, instantaneously coherent, and lagged coherent source activity, respectively, on the reconstruction of interactions in source space. These effects are generally different for different interaction measures [18].

Relatively few studies have focused on the relations between the effects of instantaneous linear mixing and the properties of the resolution operators that are used to reconstruct the source activity [12, 25, 3, 17, 24, 8]. This is perhaps surprising, because the way in which source signals are mixed is completely determined by the structure of the used resolution operator [6, 11] which is known and can therefore be exploited. For example, in [12, 3], resolution operators are exploited to design brain parcellations that minimize mixing of the reconstructed source signals and in [25, 17, 8], it is used to construct interaction measures that are insensitive to zeroth- and first-order mixing effects, whereas still being sensitive to instantaneous interactions. Indeed, classical interaction measures that do not make use of the resolution operator, such as the imaginary coherence [15], the (weighted) phase-lag index [21, 23], the imaginary phase-locking value [19], and the lagged coherence [20], are insensitive to zeroth- and first-order mixing effects, but are per construction insensitive to instantaneous interactions. It thus seems that studying the relation between mixing effects and resolution operators can provide new insights into and methods for the analysis of functional brain connectivity.

This motivates the current study, which aims to clarify some basic relations between the properties of resolution operators and the effects of linear mixing on the reconstruction of functional interactions. We focus on second-order effects because they are the least well studied and because it has recently come to light that they can cause large numbers of false positives in EEG/MEG functional connectivity analysis, even when using classical interaction measures [18]. Furthermore, we focus on second-order effects on the imaginary part of the cross-spectral function of cortical activity, since this is the simplest interaction measure that is insensitive to zeroth- and first-order mixing. The analysis of second-order effects on normalized interaction measures such as the phase-lag index and the lagged coherence is much more complicated due to their highly non-linear behavior and will be left for a future study. We do note, however, that normalization is not strictly necessary when the measures are used as test-statistics in a hypothesis test for significant interaction, because it only serves to obtain measures whose null-distribution is independent of other model parameters (e.g. the variances of the signals) but does not change the conclusion of the test.

In our analysis, we adopt a spatially continuous description of cortical activity, because it is physically the most realistic [1], because it is more general than a discrete description (they latter is a special case of the former), and because it is more natural when the aim is to obtain a basic understanding of the phenomena. Furthermore, since in EEG/MEG studies the interest is often in oscillatory activity, we model cortical activity in the frequency domain. Thus, cortical activity is modeled as a zero-mean complex-valued random stochastic field on the cortical manifold. We assume the field to be Gaussian so that it is completely described by its cross-spectral function.

The mapping from true fields to their reconstructions is modeled by a linear integral operator with a real-valued kernel, which models the composition of a linear forward operator and a linear inverse operator and is a generalization of the resolution operator to continues space [6]. The real-valuedness of the reconstruction kernel corresponds to the instantaneous nature of the forward mapping [5]. This property is crucial in the analysis of functional connectivity in source space and the classical interaction measures are insensitive to zeroth- and first-order mixing precisely because of this property.

We first characterize the effects of zeroth-, first-, and second-order mixing in terms of the crosstalk functions of the reconstruction kernel and derive alternative representations of the first- and second-order effects in terms of the symmetry properties of configurations of measurement and source locations. The representation of the second-order effects will form the basic of the subsequent analyses. In this representation, the contribution of a lagged interaction between cortical activity at a given pair of locations to the reconstructed lagged interaction between another pair of locations is proportional to a suitably defined notion of symmetry of the configuration under interchanging the measurement locations *x* and *x′*.

This representation is the generalization of that described in [17, 8] to continuous space. We use it to characterize the configurations with maximal and minimal second-order effects, to obtain a local approximation of second-order mixing effects in terms of the curvature of the cross-talk functions, to show that second-order effects are not limited to regions surroundings the measurement locations, and to establish the existence of trade-off between zeroth- and first-order effects. This trade-off is described in the form of Lagrange’s identity for pairs of cross-talk functions and relates the magnitudes of the zeroth- and second-order effects. We also provide a geometric interpretation of this identity in which the magnitude of second-order leakage between a given pair of locations is identified with the surface area of the parallelogram spanned by the two cross-talk functions and the magnitude of zeroth-order effects can be identified with their inner product. This provides a direct geometric intuition for the existence of this trade-off.

## 2 Materials and methods

### 2.1 Random neural fields

We model cortical activity in the frequency domain by a zero-mean stationary Gaussian random field on the cortical manifold Ω. The frequency coefficient of the field at a location *x* ∈ Ω is denoted by 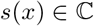 and is considered to be a random variable. Stationarity and Gaussianity together imply that the field is completely described by its cross-spectral function

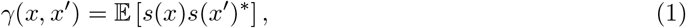

for all *x,x′* ∈ Ω, where the superscript * the complex-conjugate and 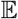 denotes expectation over temporal windows (in the case of ongoing activity) of over trials (in the case of induced activity). Note that the cross-spectral function is conjugate-symmetric, i.e. *γ*(*x′,x*) = *γ*(*x,x′*)^*^ for all *x,x′* ∈ Ω so that Re(*γ*(*x′, x*)) = Re(*γ*(*x, x′*)) and Im(*γ*(*x′, x*)) = –Im(*γ*(*x, x′*)). Also, *γ* (*x,x*) ≥ 0 is the power of the cortical activity at location *x*, which we will denote by *σ*^2^(*x*).

Following the terminology used in the field of spatial statistics [22], a neural field is called *homogeneous* if *γ*(*x,x′*) = *γ*(*d*(*x, x′*)) for all *x,x′* ∈ Ω, where *d*(*x, x′*) is a distance measure on the cortical manifold (e.g. the geodesic distance). In particular, a homogeneous field has constant power: *σ*^2^(*x*) = *σ*^2^ for all *x* ∈ Ω. A neural field is *incoherent* if *γ*(*x, x′*) = *σ*^2^(*x*)*δ*(*x* – *x′*) for all *x, x′* ∈ Ω and *coherent* if |*γ*(*x, x′*)| is constant, where the vertical bars denote the absolute value. A neural field can be represented as *s*(*x,t*) = *α*(*x, t*) exp (*iϕ*(*x, t*)), where *α*(*x, t*) and *ϕ*(*x,t*) are the associated *amplitude*- and *phase-fields*, respectively.

Eq. (1) does not allow modeling completely incoherent fields, since the cross-spectral function of such a field is given by a delta function:

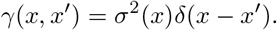

To incorporate this case, instead of Eq. (1) the cross-spectral function can be modeled as

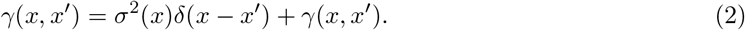

### 2.2 Linear instantaneous mixing of neural fields

When a linear inverse operator is used to reconstruct a neural field s, either based on observed electric potentials (EEG and ECoG) or based on the magnetic fluxes outside the head (MEG), the reconstructed field *s* is related to the true field by

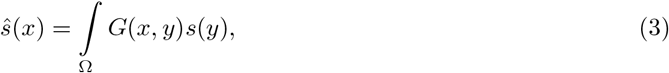

where *G*(*x, y*) is the *resolution kernel*. The mapping from *s* to *ŝ* is the concatenation of a linear forward operator and a linear inverse operator, which are left implicit here. The forward operator describes how the neural field is mapped to the sensors and in practice is obtained by numerically solving the quasi-static Maxwell equations [5, 14]. The inverse operator can be non-adaptive such as the minimum norm operator [4] or adaptive such as a beamformer [7]. The resolution kernel describes how the neural field *s* is *mixed* to obtain its reconstruction *ŝ*. Eq. (3) shows that this mixing is i.e. the reconstruction of a superposition of fields (with complex-valued coefficients) is the superposition of the reconstructed fields, and instantaneous, because the operator is real-valued.

The resolution kernel assigns to every pair of cortical locations *x, y* ∈ Ω a real number *G*(*x, y*) that determines how strong the true field at y contributes to the reconstructed field at *x*. In particular, the diagonal of the resolution kernel, i.e. the mapping (*x, x*) ↦ *G*(*x, x*) determines the gain of the reconstructed field at *x*. For *y* ≠ *x*, this can be considered to be the amount of “leakage” from *y* to *x*. In these terms, the well-known surface bias of linear inverse operators is reflected in a low gain in cortical sulci and subcortical structures and a high gain in locations that are closer to the sensors [4]. One class of resolution kernels are obtained by assuming that the leakage from *y* to *x* only depends on the (Euclidean) distance ║*x* – *y*║ between *y* and *x*. This corresponds to modeling the kernel as *G*(*x,y*) = *f*(║*x* – *y*║) for some function *f*. Typically, *f* decreases with increasing distance, e.g. *f*(║*x* – *y*║) = 1/(1 + ║*x* – *y*║) or *f*(║*x* – *y*║) = exp(– ║*x* – *y*║^2^).

When viewed as a function of *x* for fixed *y*, *G*(*x, y*) is referred to as the *point-spread function* of the kernel at *y* and when viewed as a function of *y* for fixed *x, G*(*x, y*) is referred to as *cross-talk function* of the kernel at *x*. This terminology follows that used in the discrete case [6]. If the resolution operator is symmetric, i.e. *G*(*y,x*) = *G*(*x,y*), the point-spread functions are equal to the cross-talk functions. The cross-talk functions will play a central role in this study and we will denote them by *g_x_*(*y*). Since cross-talk functions can be added and multiplied by real-valued scalars, they form an infinite-dimensional vector space over 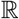. We define the following inner product on this vector space by

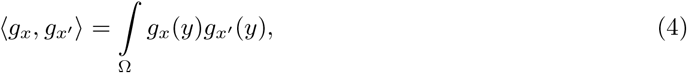

and denote the associated norm by 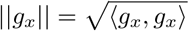. Since Ω is compact, this turns the vector space into a Hilbert space over 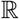.

### 2.3 Cross-spectral functions of reconstructed fields

The cross-spectral function 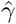 of a reconstructed random neural field is related to the cross-spectral function of the true field by

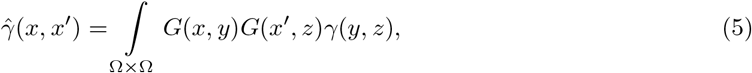

where × denotes the Cartesian product. Below, we will refer to 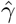 simply as the reconstructed cross-spectral function. The non-negative definiteness of the reconstructed cross-spectral function follows directly from that of the true spectrum. The reconstructed cross-spectral function can be decomposed as

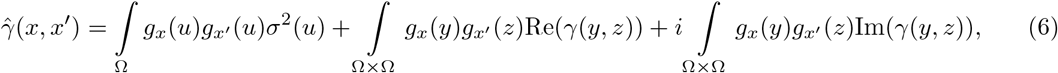

where 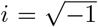 denotes the imaginary unit. We refer to the terms on the right-hand-side as zeroth-, first-, and second-order terms, respectively. Note that the zeroth-order term is independent of the interactions in the true field as described by *γ*(*y, z*) with *z* ≠ *y*, and only depends on its power *σ*^2^(*u*) at different cortical locations *u* ∈ Ω. The first- and second-order terms, on the other hand, only depend on the, respectively, instantaneous (i.e. real) and lagged (i.e. imaginary), interactions of the true field and are independent of its power. This follows from the fact that when restricted to *y* = *z*, the above integrals over Ω × Ω are zero because the diagonal of Ω × Ω is a subset of Ω × Ω with measure zero. This is the reason that the delta function was included in Eq. (2).

The above decomposition emphasizes a basic property of random neural fields and their reconstructions, which is that instantaneous interactions can only give rise to instantaneous interactions in the reconstructed field and the same holds for lagged interactions. This implies that if a reconstructed cross-spectral function has non-vanishing imaginary part, the true field must exhibit lagged interactions. It is this basic property that enables the construction of interaction measures that are insensitive to first- and second-order mixing.

If the true field is incoherent, i.e. *γ*(*x, x′*) = *σ*^2^(*x*)*δ*(*x* – *x′*), Eq. (5) only has a zeroth-order term:

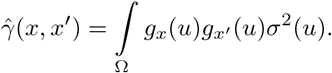

If the field is also homogeneous, i.e. *σ*^2^(*x*) = *σ*^2^, the zeroth-order term reduces to

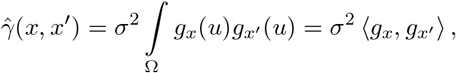

and, in particular, the reconstructed power is 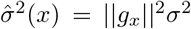. This direct relation between the zeroth-order effect of mixing and the inner product between the cross-talk functions is a well-known property of linear inverse operators. In Section 8 we will characterize the relation between zeroth- and second-order effects and the cross-talk functions and will see that this involves a generalization of the cross-product between the cross-talk functions known as the wedge product. To obtain this characterization, it will be convenient to first derive a different representation of the reconstructed cross-spectral function, which will be done in Section 5. This representation will also be used in Sections 6 and 7 to derive some other basic properties of mixing effects.

### 2.4 Basic identities for first- and second-order mixing

The conjugate symmetry of the cross-spectral function can be used to obtain formulas for the real and imaginary parts of the reconstructed cross-spectral function that simplify the analyses of mixing effects in the subsequent sections. Specifically, in Appendix A, we derive two identities that express the real/imaginary part of the reconstructed cross-spectral function in terms of the real/imaginary part of the true cross-spectral function. For *x* ≠ *x′* these identities are:

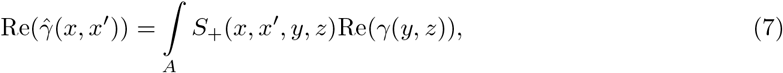

and

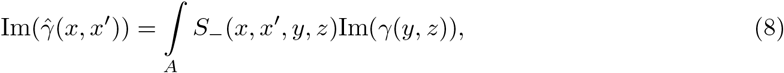

where the integral is taken over the set *A* = {(*y*_1_, *y*_2_, *z*_1_, *z*_2_)|*y*_1_ > *z*_1_}. The functions *S*_+_ and *S*_−_ are defined by

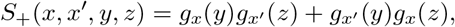

and

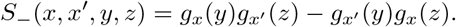

We will refer to an ordered quadruple (*x, x′, y, z*), in which *x* and *x′* are measurement locations and *y* and *z* are source locations, as a *configuration*. The squared values 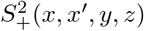 and 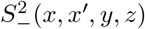 can be interpreted as measures of the lack of anti-symmetry and symmetry, respectively, of the configuration (*x, x′, y, z*), under interchanging the measurement locations *x* and *x′*. Symmetry in this context does not refer to spatial symmetry, but to the more abstract notion of invariance of an object under a class of transformations. Thus, the lack of symmetry and anti-symmetry of a configuration are directly related to the mixing effects and both are determined by the quantities *g_x_*(*y*), *g_x′_*(*z*), *g_x_*(*z*), and *g_x′_*(*y*).

Eq. (5) shows that the contribution of a true instantaneous interaction Re(*γ*(*y, z*)) between *y* and *z* to the reconstructed instantaneous interaction 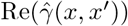 between *x* and *x′* is proportional to the lack of anti-symmetry of the configuration (*x, x′, y, z*). Likewise, Eq. (6) shows that the contribution of a true lagged interaction between *y* and *z* to the reconstructed lagged interaction between *x* and *x′* is proportional to the lack of symmetry of the configuration (*x, x′, y, z*). In other words, *S*_+_(*x, x′, y, z*)^2^ and *S*_−_(*x, x′, y, z*)^2^ are measures for the strength of, respectively, first- and second-order mixing of the true interaction between y and z into the reconstructed interaction between *x* and *x′*.

Note that *S*_+_(*x, x′, y, z*) = 0 precisely when *g_x_*(*y*)*g_x′_*(*z*) = – *g_x′_*(*y*)*g_x_*(*z*), i.e. when *g_x_*(*y*)*g_x′_*(*z*) flips sign under the transformation that interchanges *x* and *x′*. We refer to such a configuration as *antisymmetric*. Likewise, *S*_−_(*x, x′, y, z*) = 0 precisely when *g_x_*(*y*)*g_x′_*(*z*) = *g_x′_*(*y*)*g_x_*(*z*), i.e. when *g_x_*(*y*)*g_x′_*(*z*) in invariant under the transformation that interchanges *x* and *x′*. We refer to such a configuration as *symmetric*. Thus, symmetric/anti-symmetric configurations are those for which second-order/first-order mixing effects are absent. By summing 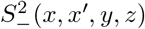 and 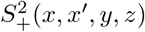 we find that they satisfy

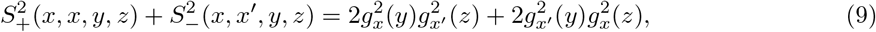

which holds for all configurations (*x, x′, y, z*). The term on the right-hand-side of Eq. (9) is a measure for the total (i.e. first- and second-order) strength of mixing of the true interaction between *y* and *z* into the reconstructed interaction between *x* and *x′* and hence shows that, given the total mixing strength, there is a trade-off between the strength of first- and second-order mixing. In particular, given the total mixing strength, symmetric configurations have maximal first-order mixing effects and anti-symmetric configurations have maximal second-order mixing effects.

## 3 Results

### 3.1 Configurations with maximal and minimal mixing effects

To illustrate the use of the basic identities (Eqs. (7) and (8)) we provide characterizations of the configurations that are maximally and minimally sensitive to first- and second-order mixing effects. These characterizations will be in terms of the configurations’ geometry, i.e. in terms of the relative positions of the measurement and source locations. Because these positions determine the symmetry of the configuration only indirectly via the cross-talk functions, to relate the symmetry to the geometry of a configuration, we need to make some mild assumptions about the cross-talk functions. We assume that for every location *x*, (i) *g_x_* is maximal in *x*, (ii) *g_x_* decreases with increasing Euclidean distance from *x*, and (iii) *g_x_* is non-negative, i.e. *g_x_* ≥ 0.

We ask which configurations (*x, x′, y, z*) minimize/maximize *S*_−_(*x, x′, y, z*)^2^ and *S*_+_(*x, x′, y, z*)^2^. The only non-trivial case is for which configurations *S*_−_(*x, x′, y, z*)^2^ is minimized and will be discussed last. First consider for which configurations

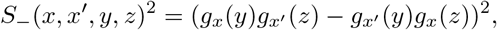

is maximized. This is the case if *g_x_*(*y*)*g_x′_*(*z*) is maximal and *g_x′_*(*y*)*g_x_*(*z*) is minimal (or the other way around). Now, *g_x_*(*y*)*g_x_*(*z*) is maximal if *y* = *x* and *z* = *x′* (assumption (i)), in which case *g_x′_*(*y*)*g_x_*(*z*) reduces to *g_x′_*(*x*)*g_x_*(*x′*), which is minimal if *x* and *x′* are far apart (assumption (ii)). Thus, the configurations that are maximally sensitive to second-order mixing effects are those for which the source locations coincide with the measurement locations and for which the measurement locations are far apart. Now consider for which configurations

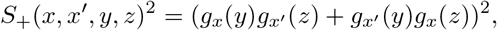

is maximized. This is the case if both terms *g_x_*(*y*)*g_x′_*(*z*) and *g_x′_*(*y*)*g_x_*(*z*) are maximal. The first term is maximal if *y* = *x* and *z* = *x′* (assumption (i)). This reduces the second term to *g_x′_*(*x*)*g_x_*(*x′*), which is maximal if *x* = *x′* (assumption (i)). Thus, the configurations that are maximally sensitive to first-order mixing effects are those in which both measurement and source locations coincide. From the above expression for *S*_+_(*x, x′, y, z*)^2^ it is also immediately clear that *S*_+_(*x, x′, y, z*)^2^ is minimal if both terms *g_x_*(*y*)*g_x′_*(*z*) and *g_x′_*(*y*)*g_x_*(*z*) are zero, which is the case if one of the cross-talk functions in each term is zero, for which there are four possibilities. Two of these are that both sources are sufficiently far from both measurement locations (assumption (i)).

We are left with the question for which configurations is *S*_−_(*x, x′, y, z*)^2^ is minimal, or equivalently, for which configurations *g_x_*(*y*)*g_x_*(*z*) = *g_x′_*(*y*)*g_x_*(*z*). There is no general answer to this question, because it depends on the particular form of the cross-talk functions *g_x_* and *g_x′_*. Consider the special case

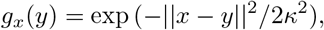

were ║*x* – *y*║ denotes the Euclidean distance between *x* and *y* and *κ* is the characteristic scale of *g_x_*. The above condition then takes the form

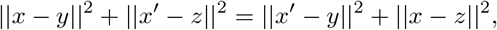

which is equivalent to 〈*x* – *x′, y* – *z*〉 = 0, where the brackets denote the dot product in 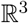. This shows that sensitivity to lagged interactions is zero precisely when the line through the measurement locations *x* and *x′* and the line through the source locations *y* an *z* are perpendicular. In particular, for fixed source locations *y* and *z*, the measurement locations for which sensitivity to lagged interactions is zero form a two-dimensional plane in 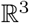. If the configuration is confined to a two-dimensional plane, they form a line in 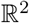, and in the configuration is confined to a line, sensitivity to lagged interactions is never zero.

In general, characterizations of the symmetric configurations are more complicated than in the above special case. For example, if *g_x_*(*y*) is a rational function (i.e. a quotient of polynomials) in ║*x* – *y*║^2^, the planes will be replaced by curved surfaces. An example is

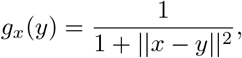

in which case the symmetric configurations are characterized by the condition

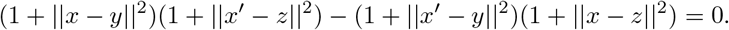

For fixed source locations *y* and *z*, the solutions of this equation form a two-dimensional surface in 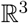.

### 3.2 Mixing effects in the vicinity of the measurement locations

In the previous section we established that the configurations that are maximally sensitive to second-order mixing effects are those for which the source locations coincide with the measurement locations and for which the measurement locations are far apart. We now consider the case that the sources are located in the vicinity of the measurement locations (and the latter are far apart). This case was explored extensively using numerical simulations in [18]. Specifically, we relate *S*_−_(*x, x′, y, z*) to the geometric properties of the cross-talk functions *g_x_* and *g_x′_* in the vicinity of *x* and *x′*, respectively.

If *x* and *x′* are far apart, we neglect the term *g_x′_*(*x*)*g_x_*(*x′*) in *S*_−_(*x, x′, y, z*) and approximate *g_x_*(*y*) and *g_x′_*(*z*) by second-order Taylor series in *x* and *x′*, respectively. For *y* close to *x* and *z* close to *x′*, that *S*_−_(*x, x’, y, z*) can then be approximated as

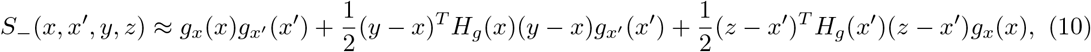

where *H_g_*(*x*) and *H_g_*(*x′*) denote the Hessian matrices of *g_x_* at *x* and of *g_x′_* at *x′*, respectively (see Appendix B). Thus, the (*i, j*)-th entry of *H_g_*(*x*) is given by the second-order partial derivative of *g_x_* to *x_i_* and *x_j_* at *x*:

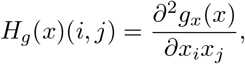

where *∂_ij_g_x_*(*x*) denotes the second-order partial derivative of *g_x_* to the *i*-th and *j*-th coordinates of *x* = (*x*_1_, *x*_2_, *x*_3_) at point 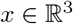. This approximation is also valid for *S*_+_(*x, x′, y, z*). Eq. (9) shows that the local structure of signal leakage is determined by the Hessian matrices of the cross-talk functions at the two measurement locations and that the effects are additive. Assumptions (i) and (ii) imply that *H_g_*(*x*) and *H_g_*(*x′*) are negative definite. Combining this with assumption (iii) we conclude that the second and third term on the right-hand-side of Eq. (9) are negative. From this we can conclude that leakage is maximal at the measurement locations and decreases in the neighborhood of the measurement locations.

As a special case, suppose that both Hessian matrices are proportional to the 3 × 3 identity matrix, with proportionality constant –*ξ* for some *ξ* > 0, i.e. *H_g_*(*x*) = (*x′*) = – *ξI*_3_, where *I*_3_ denotes the 3 × 3 identity matrix. Then Eq. (9) reduces to

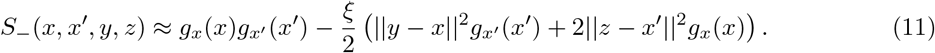

The Gaussian curvature *K* of *g_x_* at *x* and of *g_x′_* at *x′* is given by the determinant of the respective Hessian matrices, which equals *ξ*^3^, we see that the local leakage effect is proportional to *K*^1/3^. If, in addition, *g_x_*(*x*) = *g_x′_*(*x′*) = 1 and ║*y* – *x*║ = ║*z* – *x*| = *δ*, we obtain the following approximation:

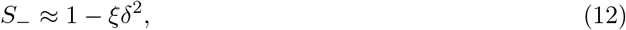

which is also valid for *S*_+_. It shows that, in the vicinity of the measurement locations, leakage decreases linearly with *ξ* and quadratically with distance. As an example, let *g_u_*(*v*) = exp(– ║*u* – *v*║^2^/2*κ*^2^), where *κ* > 0 is the characteristic width of *g_u_*. Thus, large values of *κ* correspond to a low spatial resolution. Its Hessian is proportional to the 3 × 3 identity matrix with *ξ* = 1/*κ*^2^ so that *S*_−_ ≈ 1 – *δ*^2^/*κ*^2^.

### 3.3 Relation between total zeroth- and second-order mixing effects

In Section 2.4 we showed that, for a given configuration of measurement and source locations, and given the total mixing strength, there is a trade-off between first- and second-order mixing effects. We now show that a related trade-off exists when integrated over all true interactions.

As a measure for the total strength of zeroth-order mixing effects in the reconstruction of the interaction between *x* and *x′* we take the squared inner product of the cross-talk functions at *x* and *x′*:

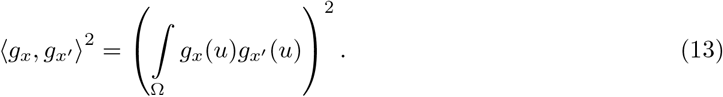

Note that zeroth-order effects are absent precisely when *g_x_* and *g_x_* are orthogonal. As a measure for the total strength of second-order mixing we take the quantity ║*g_x_* ∧ *g_x′_*║^2^, which we define by

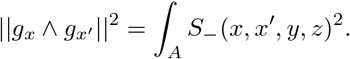

The notation ║*g_x_* ∧ *g_x′_*║ will be explained in Section 3.5 when we discuss the special case of finitely many point-sources. We first discuss the following relation, which is derived in Appendix B:

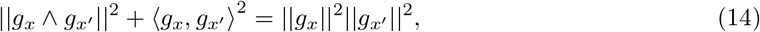

where ║*g_x_*║ denotes the norm of *g_x_* (see Section 2.2). It is a continuous version of Lagrange’s identity and shows that, given the norms ║*g_x_*║ and ║*g_x′_*║ of the cross-talk functions, there is a trade-off between the total zeroth- and second-order mixing effects to the reconstruction of the interaction between *x* and *x′*.

Note that, given ║*g_x_*║ and ║*g_x′_*║, if *g_x_* and *g_x′_* are orthogonal, zeroth-order effects are absent and second-order effects are maximal and if *g_x_* and *g_x′_* are linearly dependent, second-order effects are absent and zeroth-order effects are maximal. By dividing both sides of Eq. (14) by ║*g_x_*║^2^║*g_x′_*║^2^, we obtain the following relation

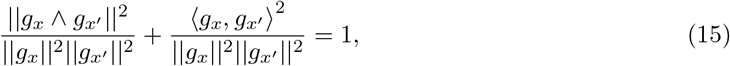

between the normalized measures of the strengths of zeroth- and second-order mixing effects. This relation can also be written as

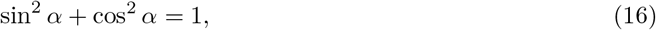

where *α* denotes the angle between *g_x_* and *g_x′_* and clearly shows the trade-off between the relative strengths of zeroth- and second-order mixing effects.

Lastly, we provide a geometric interpretation of the trade-off in Eq. (14) in terms of the parallelogram spanned by *g_x_* and *g_x′_*. For two finite-dimensional vectors *v* and *v′*, the area of the parallelogram spanned by v and v’ equals the square root of the determinant of their Gram matrix:

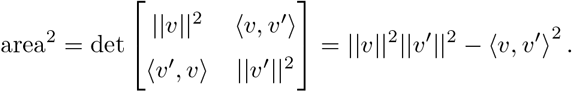

This can be generalized to infinite-dimensional Hilbert spaces, which allows to define the area of the parallelogram spanned by two cross-talk functions. Thus, the area of the parallelogram spanned by two cross-talk functions *g_x_* and *g_x′_* can be defined as

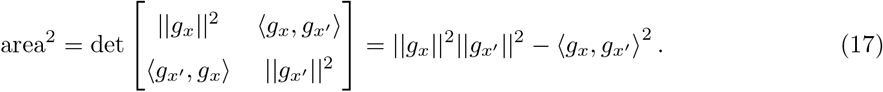

Note that the area is always non-negative because the Gramm matrix is non-negative definite. Comparing Eq. (17) with Eq. (14) we conclude that

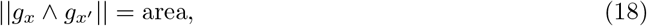

which shows that the total strength of second-order mixing to the reconstructed interaction between *x* and *x′* can be interpreted as the area of the parallelogram spanned by the cross-talk functions *g_x_* and *g_x′_*.

### 3.4 The discrete case

In this section we consider the special case that the neural field comprises *N* point-sources at locations *x*_1_,…, *x_N_* ∈ Ω. This largely amounts to replacing integrals by sums, but will also provide some geometric insight into the structure of mixing effects. The neural field in this case takes the following form:

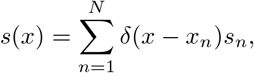

where *s_n_* is the Fourier coefficient of the *n*-th source. Let 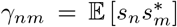 be the cross-spectrum between the *n*-th and *m*-th source. The cross-spectrum *γ*(*x, x′*) between measurement locations *x* and *x′* now becomes

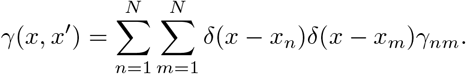

Furthermore, the reconstructed field takes the form

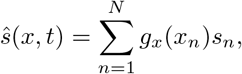

where *g_x_* is the cross-talk function at *x*. We assume that the measurement locations do not coincide with the source locations so that the reconstructed field is entirely spurious. Note that the cross-talk functions *g_x_* and *g_x′_* are now *N*-dimensional vectors and that *g_x_*(*x_n_*) is the *n*-th coordinate of *g_x_*.

The cross-spectrum of the reconstructed field is

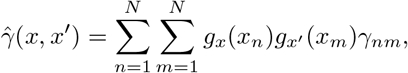

which can be decomposed as

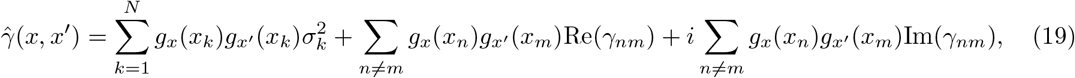

where 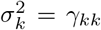 is the power of the *k*-th source. Eq. (19) is the discrete version of Eq. (6). Note that the first term (zeroth-order mixing) is independent of the interaction structure of the sources and only depends on their power, whereas the second and third terms (first- and second-order mixing, respectively) only depend on the, respectively, instantaneous and lagged, interaction structure and are independent of power. The basic identities form Section 2.4 take the form

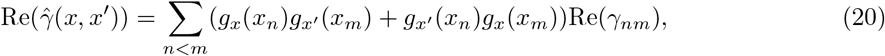

and

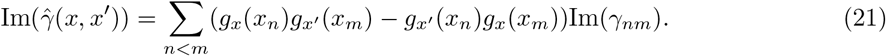

The term ║*g_x_* ∧ *g_x′_*║^2^ reduces to

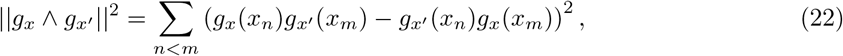

hence the notation *g_x_* ∧ *g_x′_* now gets meaning because it equals the *wedge product* between the vectors *g_x_* and *g_x′_*, which is defined as

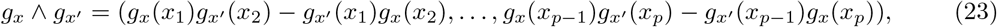

and is a vector of dimension *N*(*N* – 1)/2. Strictly speaking, the wedge product is not a vector at all, but an oriented plane spanned by *g_x_* and *g_x′_*. It is a generalization of the cross product to higher dimensional vectors: In the special case *N* = 3, *g_x′_* = (*a, b, c*) and *g_x_* = (*d, e, f*) are vectors in 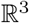 and (after permutation of its entries and signs) their wedge product reduces to

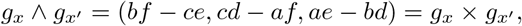

where *g_x_* × *g_x′_* denotes the cross-product between *g_x_* and *g_x′_* and, as such, satisfies the same properties, e.g. *g_x_* ∧ *g_x_* = 0 and *g_x′_* ∧ *g_x_* = –*g_x_* ∧ *g_x′_*. In fact, the wedge product is uniquely determined by these axioms. Lagrange’s identity for cross-talk functions hence reduces to

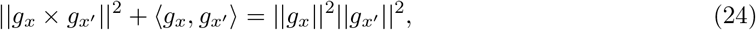

and in the normalized case this reduces to Eq. (16).

## Discussion

In this study we established several basic properties regarding the effects of instantaneous linear mixing on the reconstructed cross-spectra of random neural fields. Although these properties are rather superficial from a mathematical point of view, they do provide some insight into aspects of signal mixing that are relevant to experimentalists working with EEG or MEG data. For instance, one of the results is that second-order mixing effects are most severe when the measurement locations are far apart and the sources are located in the vicinity of the measurement locations. The result provides some formal understanding of the large number of false positive interactions in the vicinity of the measurement locations as observed via numerical simulations [18].

Although mixing effects are usually studied within a discrete framework by discretizing the source space, we used continuous kernels and random neural fields on cortical manifolds, because it allows for a more natural description of some of the effects, e.g. the relation between mixing effects and the curvature of the point-spread functions. Another reason for adopting this framework is that macroscopic cortical activity is a spatiotemporal phenomenon exhibiting properties such as traveling waves, which are more naturally studied within such a framework. For instance, a description of mixing effects in the spatial frequency domain is readily obtained from the continuous description used in this study by taking the spatial Fourier transforms of the neural fields and the resolution kernels [11].

An obvious next question is how the coherences (i.e. normalized cross-spectra) of the true and reconstructed fields are related. Although the decomposition into zeroth-, first-, and second-order effects (see Eq. (6)) is still valid, the coherence is a non-linear function of the different terms and this considerably complicates the analysis. One faces similar difficulties when analyzing the relationship between non-linear properties of the true and reconstructed fields, for example their phase- or amplitude-dynamics. For example, in [9] forward simulations are used to explore the highly non-linear relation between true and observed phase-fields in the context of local field potential recordings. One of the effects that could be analyzed mathematically is phase-contraction, which refers to the fact that the phase-difference between reconstructed signals is typically smaller than that between the true signals, which leads, for example, to overestimation of propagation speeds of neural activity.

The resolution operator was modeled by a linear integral operator with general real-valued kernel and some simple choices for the kernel were considered (e.g. a Gaussian kernel). Although this allowed to clarify some basic aspects of second-order signal leakage, the study of specific effects requires making *ad hoc* choices. For example, effects of source depth can be incorporated by suitably parametrizing the kernel, e.g. by multiplying it with a positive constant < 1 and increasing its spatial width. These choices, however, are not derived from first principles. To arrive at a more fundamental formalism, the forward and inverse operators that make up the resolution operator should be modelled explicitly. For MEG, the forward operator is given by the Ampere-Laplace law, and for EEG and ECoG, the forward operator is given by the integral form of Poisson’s equation, both of which are linear integral operators [5]. Effects of source depth, unknown dipole orientation, etc. can then be studied from first principles without the need for *ad hoc* choices. Since the brain is modeled as a spatial continuum, it does, however, require the use of inverse operators that map sensor activity into a Hilbert space of brain activity (in contrast to a finite-dimensional vector space). In contrast to the field of electromagnetic brain imaging [4], such continuous formulations of inverse methods are standard in most other fields, e.g. acoustic scattering, optical tomography, and seismology, and enable rigorous mathematical analysis (for instance, see [26]).

## Appendices

## Appendix A: Derivation of the basic identities

The above formulas can be derived in the following way. Let 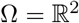 so that for *x*, 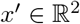:

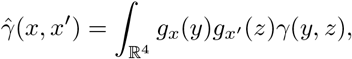

where *y* = (*y*_1_, *y*_2_) and *z* = (*z*_1_, *z*_2_) are in 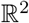. Define subsets of 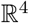 by *A*^+^ = {(*y*_1_, *y*_2_, *z*_1_, *z*_2_)|*y*_1_ > *z*_1_} and *A*^−^ = {(*y*_1_, *y*_2_, *z*_1_, *z*_2_)|*y*_1_ < *z*_1_} and write

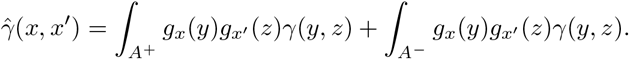

Now define 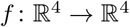 by *f*(*x, y*) = (*y, x*) and note that *f* induces a diffeomorphism between *A*^+^ and *A*^−^ with (absolute value of) determinant 1 so that we can write the second term of the above equation as:

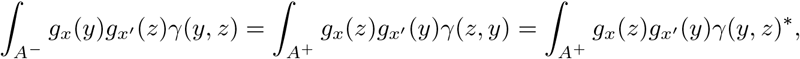

where in the last step we have used the conjugate symmetry of *γ*, i.e. *γ*(*z, y*) = *γ*(*y, z*)^*^ for all *x*, 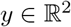. We can thus rewrite 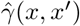 as an integral over *A*^+^:

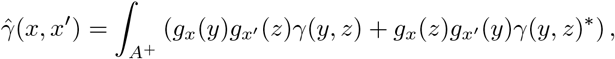

and its real and imaginary parts therefore are

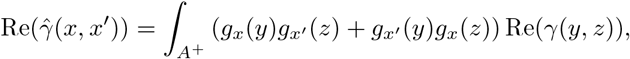

and

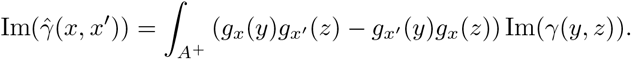

## Appendix B: Local approximation of second-order leakage

We consider the configuration (*x, x′, y, z*), where *x* and *x′* are measurement locations and *y* and *z* are source locations and assume that *y* is close to *x* and *z* is close to *x′*. We also assume that, for all locations *v*, the cross-talk function *g_v_*(*u*) is maximal at *u* = *v* and decreases with increasing Euclidean distance between *u* and *v*. Under these assumptions, *g_x′_*(*y*)*g_x_*(*z*) ≈ 0 so that *S*_−_(*x, x′, y, z*) and *S*_+_(*x, x′, y, z*) are approximately equal to *g_x_*(*y*)*g_x′_*(*z*). Since *y* is close to *x* and *z* is close to *x′*, *g_x_*(*y*) and *g_x′_*(*z*) can be approximated by second-order Taylor series around *x* and *x′*, respectively:

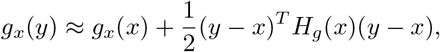

and

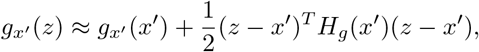

where *H_g_*(*x*) and *H_g_*(*x′*) denote the Hessian matrices of *g_x_* at *x* and of *g_x_* at *x′*, respectively. Thus, the (*i,j*)-th entries of *H_g_*(*x*) and *H_g_*(*x′*) are equal to *∂*^2^*g_x_*(*x*)/*∂x_i_x_j_*, and *∂*^2^*g_x′_*(*x′*)/*∂x′_i_x′_j_*, respectively. The first-order terms in the Taylor series of *g_x_* and *g_x′_* vanish due to the assumptions that *g_x_* and *g_x′_* are maximal at *x* and *x′*, respectively. Combining the approximations for *g_x_*(*y*) and *g_x_*(*z*) yields the following approximation for *S*_−_(*x, x′, y, z*):

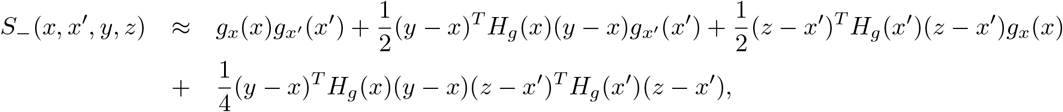

which is also valid for *S*_+_(*x, x′, y, z*). If *y* is sufficiently close to *x* and *z* is sufficiently close to *x′*, the last term on the right-hand side will be much smaller than the second- and the third terms so that we can further approximate *S*_+_(*x, x′, y, z*) as

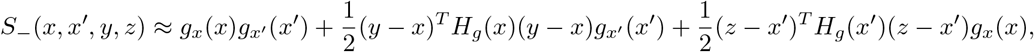

which is also valid for *S*_+_(*x, x′, y, z*).

## Appendix C: Derivation of Lagrange’s identity

To prove Lagrange’s identity, we again use the diffeomorphism *g* between *A*^+^ and *A*^−^ to write

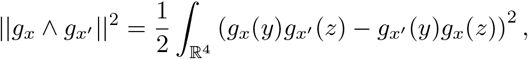

so that

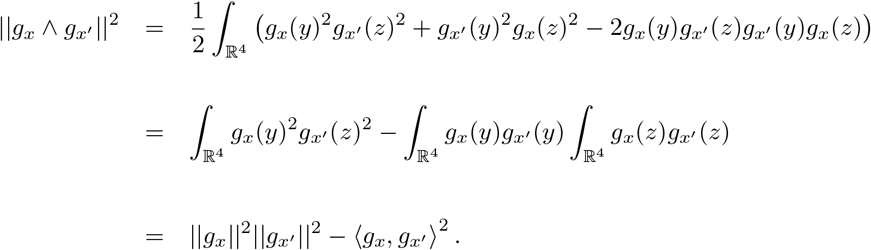

## Acknowledgments

Rikkert Hindriks was funded by NWO-Wiskundeclusters grant nr. 613.009.105.

